# Modeling Brain Connectivity Dynamics in Functional Magnetic Resonance Imaging via Particle Filtering

**DOI:** 10.1101/2021.01.19.427249

**Authors:** Pierfrancesco Ambrosi, Mauro Costagli, Ercan E Kuruoğlu, Laura Biagi, Guido Buonincontri, Michela Tosetti

## Abstract

Interest in the studying of functional connections in the brain has grown considerably in the last decades, as many studies have pointed out that these interactions can play a role as markers of neurological diseases. Most studies in this field treat the brain network as a system of connections stationary in time, but dynamic features of brain connectivity can provide useful information, both on physiology and pathological conditions of the brain. In this paper, we propose the application of a computational methodology, named Particle Filter (PF), to study non-stationarities in brain connectivity in functional Magnetic Resonance Imaging (fMRI). The PF algorithm estimates time-varying hidden parameters of a first-order linear time-varying Vector Autoregressive model (VAR) through a Sequential Monte Carlo strategy. On simulated time series, the PF approach effectively detected and enabled to follow time-varying hidden parameters and it captured causal relationships among signals. The method was also applied to real fMRI data, acquired in presence of periodic tactile or visual stimulations, in different sessions. On these data, the PF estimates were consistent with current knowledge on brain functioning. Most importantly, the approach enabled to detect statistically significant modulations in the cause-effect relationship between brain areas, which correlated with the underlying visual stimulation pattern presented during the acquisition.

## 1. INTRODUCTION

The understanding of brain functioning is linked to the study of the dynamic interaction among anatomically segregated brain areas. These interactions are labeled functional and effective connectivity and refer to distinct ways of considering connections among brain region. While complementary to structural connectivity, which describes anatomical connections between brain regions [1], they concern functional connections that are not necessarily achieved through a direct anatomical link between brain areas. Functional connectivity regards connections as statistical codependencies, and consequently it is a non-directional and model-free description of the brain network. On the contrary, effective connectivity defines the temporal relationship and causal influences in the brain in a given network model [2].

Functional Magnetic Resonance Imaging (fMRI) is frequently employed in brain connectivity studies, given its noninvasiveness and satisfactory spatiotemporal resolution, both in physiology and pathology (e.g. Alzheimer’s disease [3]–[5], schizophrenia [6] and Major Depression Disorder [7]). From brain connectivity studies it emerged that brain dynamics, in particular effective connectivity, may provide a biological marker for specific brain disease and a tool for monitoring responses to treatments of these pathologies [8]–[11].

Granger Causality (GC) and Dynamic Causal Modeling (DCM) are methods to investigate effective connectivity. GC is present when knowledge on temporal evolution of the signal in a certain brain region *A* improves the predictability of another brain region *B* [12], [13]. This approach is based on the evaluation of a linear codependence among time series, and it is therefore limited to a stationary framework or needs a sliding-window approach to address time-varying coupling between regions, which has limitations [14]. Differently, in DCM the predicted relationship between neural activity and observed fMRI signal needs to be specified in a pre-determined model, hence requiring previous knowledge about the timing and effect on signals of the connectivity modulation [15].

The Sequential Monte Carlo (SMC) methodology [16] is crucially different from these two strategies. SMC approaches estimate the hidden states of a dynamic system with only partial and noisy observations, without further assumptions on the presence of variations in connectivity. A specific SMC methodology called Particle Filter (PF) employs discrete sampling to approximate probability density functions and it updates the posteriors with the arrival of new samples.

The SMC algorithm proposed here was recently developed by Ancherbak et al. [17], originally for time-varying gene network modeling. We adapted it for the study of brain connectivity using fMRI data and the feasibility and behaviour of the proposed approach has been studied on synthetic data mimicking fMRI time-series. When applied to real fMRI datasets, results were compared to correlation between delayed time series, considered as a proxy measure for stationary effective connectivity. Two different experimental paradigms were tested, one during periodic tactile stimulation and one during visual periodic stimulation. Two different experimental paradigms were tested: the first one, whose preliminary results were presented in abstract form [18], involved tactile stimulation during an fMRI acquisition with temporal resolution of 2s. The second experiment employed a periodic visual stimulation with significantly improved time resolution of 0.8s.

## II. METHODOLOGY

### A. Model and algorithm

Particle filter [17], [19]–[22] is a sequential Monte Carlo methodology based on the Bayes theorem on conditional probability. Particle filters estimate the probability distributions of hidden variables of interest, modeled according to a hypothesized *state-space equation*. The probability density function (*pdf*) of the hidden variables is allowed to be time-varying and is therefore sequentially updated when new data become available. Such probability distribution is estimated from the data, modeled according to a hypothesized *observation equation*. In brain connectivity studies based on fMRI data, the relationship among the time-series of *R* different brain Regions of Interest (ROIs) x_*t*_ = {*x*_1_(*t*)*, … , x_R_*(*t*)} can be modeled as a first order linear Vector Autoregressive (VAR) model [12], [21], [23]–[25] as:

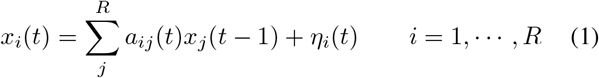

or in matrix notation:

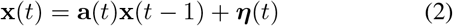

where

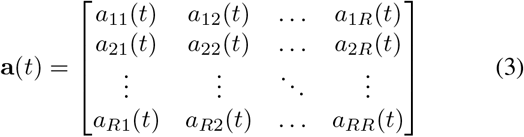

which is employed as the observation equation describing the relationship between the observations x(*t*) at time *t* and those at time *t* − 1 (that is, x(*t* − 1)); ****η****(*t*) is the vector of observation noise; the matrix of hidden parameters of interest a(*t*) represents the causal influence exerted between different areas. In particular, it can be assumed that elements of a(*t*) are allowed to be time-varying:

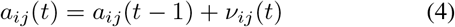

where *a_ij_*(*t*) is the *ij*-th element of the matrix a(*t*), describing the influence of the *j*-th region over the *i*-th region, and *ν_ij_*(*t*) is the process noise (innovation) term.

The PF algorithm evolves from an initial probability distribution for *a_ij_*(*t* − 1), which we chose to be uniform at *t* = 1, and through equation (4) it generates new possible values for *a_ij_*(*t*); then, with equation (2), the PF algorithm generates predicted values of the observations at time *t*. The desired probability density function of the parameters of interest a(*t*) can be estimated via Bayes theorem as follows:

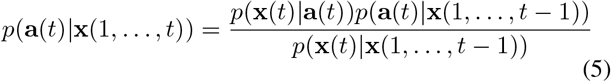

and with the assumption of Gaussian noise we have

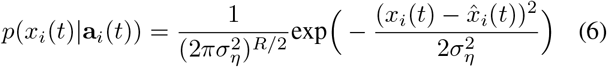

where 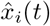 are the data estimated through equation (1) at time *t* for the *i*-th ROI and a_*i*_(*t*) = {*a_i_*_1_*, … , a_iR_*} is the vector of hidden variables associated with the *i*-th ROI at time *t*, that is, the *i*-th row of matrix a_*t*_.

In most applications, equation (5) cannot be solved analytically [26], but it can be computed through the Sequential Monte Carlo sampling scheme, which consists in representing the pdf *p*(a(*t*)|x(1*, … , t*)) as a discrete set of N weighted samples called *particles*:

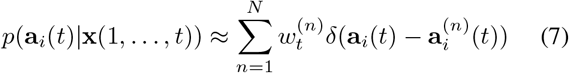

where 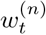 is the weight associated to the *n*-th particle vector 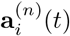 for the *i*-th row of matrix a(*t*) at time *t*. The Sequential Importance Sampling (SIS) [26] methodology provides a strategy to compute the weights. It has been shown [20] that the weights can be sequentially updated as follows:

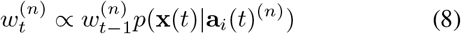

where the proportionality takes into account normalization factors. With this approach, at each time instant *t* we have a sample set 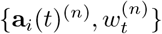 for *n* = 1*, … , N* and for *i* = 1*, … , R* which can be used to estimate the pdf of the parameters and to infer information about the network. However, after some iterations, most of the particles will have a very low statistical weight, resulting in a lower exploration efficiency of the algorithm. To overcome this typical problem of sequential Monte Carlo methodologies, a step called Resampling is performed. The number of effective particles was defined in [27] as

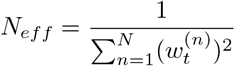

If *N_eff_* is below a certain arbitrary threshold the Resampling is performed: particles with weight below a certain threshold are substituted by copies of particles with sufficiently high weights and to each of the new particles set is assigned the same weight 1*/N*. This results in a more effective exploration of the solution space, because only statistically relevant particles remain after this step.

To sum up, the resulting algorithm can be schematically expressed as in Table I. In our implementation the procedure is repeated *N_r_* = 100 times, all independent from each other, to provide a better exploration of the solution space, and resampling was performed when *N_eff_ <* 30% of the total number of particles. The final outputs of the algorithm are the a_*t*_ computed as the average of the *N_r_* repetitions. The algorithm was implemented in MATLAB (Mathworks, Natick, MA, U.S.A.) R2017b.

**Table 1.** Schematic description of the PF algorithm.

|  |  |
| --- | --- |
| Input the BOLD fMRI data series | $\mathbf{x}_t$ |
| Set the number of particles | $N$ (=2000 in our case) |
| Set starting point for coefficients values | $\mathbf{a}_{t=0} = 0$ |
| Start PF | for $t = 1 : T$ |
| updating step | generate $N$ particles from previous coefficients' values through<br>$a_{ij}(t) = a_{ij}(t-1) + \nu_{ij}(t)$ |
| estimation step | predict the values of the observations at time $t$ from values at time $t-1$ with<br>$\mathbf{x}(t) = \mathbf{a}(t)\mathbf{x}(t-1) + \boldsymbol{\eta}(t)$<br>compute the likelihood between predicted values and observed values with (6)<br>normalize the weights and resample |
| End PF | end for on $t$ |

### B. Synthetic data

To validate the proposed approach, two different synthetic networks were used.

- One network with *R* = 6 nodes, each with *T* = 100 time points, stationary coefficients generated with the MATLAB function *varm()* with a Signal-to-Noise Ratio (SNR) set to either *∞* (*σ_η_* = 0, ideal case) or 6dB.
- Another network with *R* = 2 and *T* = 250 was used to assess the PF capability to capture time-varying hidden parameters. In this case, *a_ij_* coefficients were zero except for coefficient *a*_21_, whose value switched from 1 to 1 with a period of 125 time points. The SNR was 10dB.

The first synthetic dataset allowed us to verify the reliability of the results of the PF and to decide the optimal values of the parameters; the second one allowed us to address the capability of the PF to track variations through time of the AR coefficients.

### C. Real fMRI data

The proposed approach was also retrospectively applied to real fMRI data acquired on healthy volunteers in two different experimental set-ups, acquired at isotropic spatial resolution of (1.5 mm)^3^ on a 7T MRI system:

- **Motor Task**: Time-series consisting of 240 time points with a temporal resolution of 2*s* were acquired on two subjects. During acquisition, the subjects’ thumb- and index-fingertips were stimulated via a pneumatic device (Linari Engineering, Pisa, Italy). The subjects’ task was to move the finger whenever it was stimulated. Four ROIs were studied covering primary somatosensory (S1), primary motor (M1), supplementary motor (SM) and parietal (P) cortices. All ROIs consisted in four voxels and were manually drawn on each subject on one slice only, to avoid potential slice timing confounds (Fig. 1). The resultant time-series of each ROI were obtained by averaging the four time-series of individual voxels.
- **Visual Task**: Time-series consisting of either 300 or 600 time points with a temporal resolution of 0.8*s* were acquired on four trials. Subjects underwent a periodic visual stimulation alternating between black and white dots moving along spiral trajectories over a gray background (stimulation ON) and presentation of the gray background alone (stimulation OFF). Four ROIs were studied covering the Lateral Geniculate Nucleus (LGN), the Middle temporal cortex (MT), the Primary Visual area (V1) and one control ROI in the tempo-parietal cortex (CTRL). Four voxels wide ROIs were manually drawn on each subject on one slice only.

**Fig. 1:**
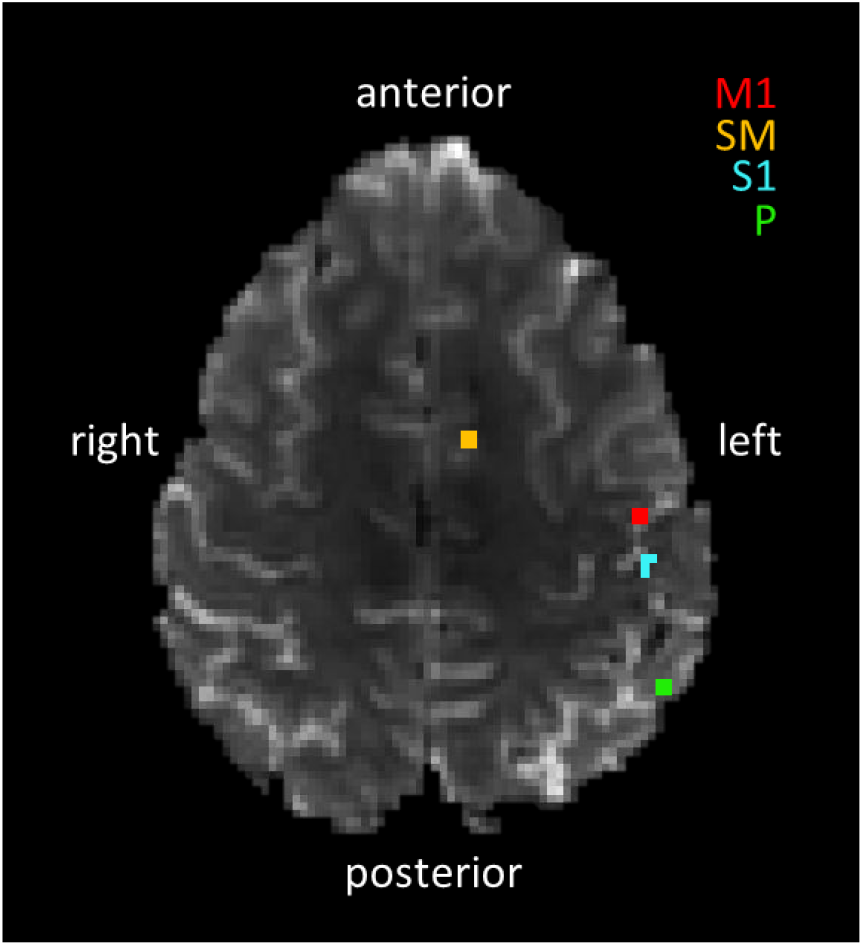
ROIs drawn on one representative subject, representing primary somatosensory (S1), primary motor (M1), supplementary motor (SM) and parietal (P) cortices.

The optimal order of the autoregressive model describing the time series was 1, as estimated by the Schwartz criterion [12].

The particle filter was applied not only to the original time series but also to the same fMRI data shuffled in time. This test allowed to address the effective dependency of the results from the temporal order of the data, i.e. to address causal dependency through time in the data. Subsequently, the *a_ij_* coefficients estimated by particle filtering were compared to the delayed correlation (DC) *c_ij_* between signals *x_i_*(*t*) and *x_j_*(*t* − 1) which reflect the time-invariant causal influence exerted by network node (ROI) *j* over node *i*. Furthermore, the results of the PF were also compared to coefficients estimated in a stationary framework fitting a four-node first order linear AR model to the data via maximum likelihood.

With a t-test between coefficient values in the presence and in the absence of stimulation we searched statistically significant (p-value *<* 0.05) changes in connectivity following the underlying stimulation pattern. Also, as a control analysis, a second t-test was performed, simulating a different stimulation pattern from the actual one, which allowed to exclude variations deriving from spurious fluctuations of the results, as explained in Fig. 2.

**Fig. 2:**
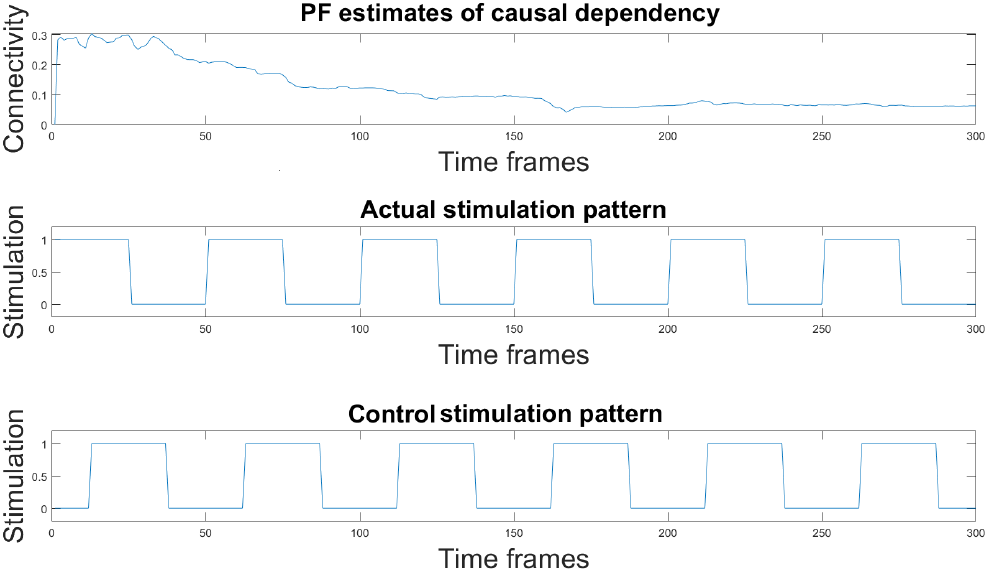
The plot on top shows an example of PF estimates of causal dependency between time-series. Plot in the center shows the actual stimulation pattern as a square wave, where the presence and the absence of stimulation are represented by 1 and 0 respectively. The bottom plot shows a time shifted stimulation pattern, which has a half-period offset from the actual stimulation pattern. The t-test was run comparing connectivity values in correspondence of ones (stimulation ON) and zeros (stimulation OFF) in both cases. The second test allowed to exclude spurious changes in connectivity values not due to the stimulation.

## III. EXPERIMENTAL RESULTS AND DISCUSSIONS

### A. Synthetic Data

Scatter plots in Fig. 3 demonstrate that the causality coefficients estimated by PF in a stationary network satisfactorily correlate with the true coefficients, both in the noiseless synthetic dataset (Pearson’s *ρ* = 0.96) and in the noisy scenario with SNR = 6dB (Pearsons’ *ρ* = 0.59). The case of a network with one time-varying coefficient is shown in Fig. 4. The PF tracks the changes of the non-stationary coefficient *a*_21_, although the estimated values do not immediately follow the abrupt changes between 1 and *−*1 and viceversa. All the other coefficients are correctly estimated to be close to the nominal null value.

**Fig. 3:**
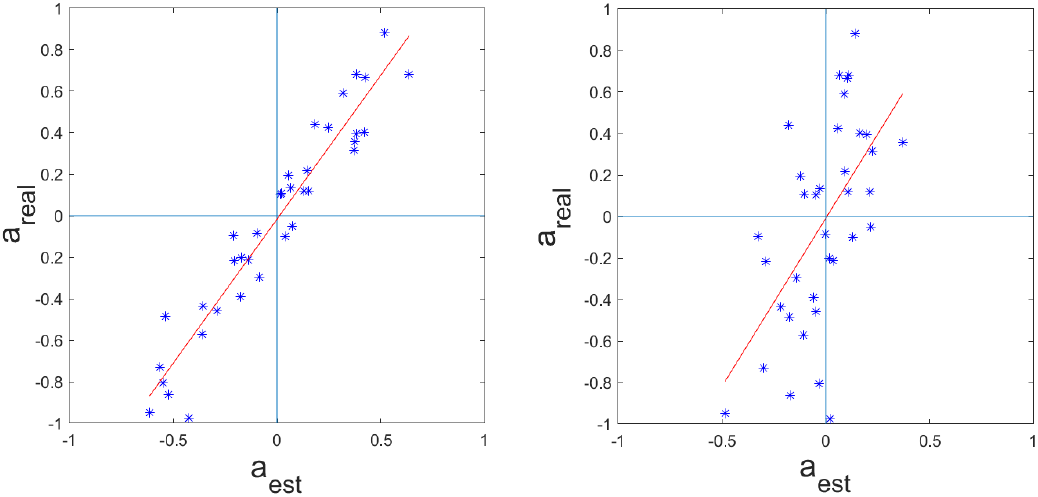
Scatter plots that relate PF estimates (x axis) and true values (y axis) of the autoregressive model for a 6-node network with 100 time samples, in the absence of noise (left) and with SNR = 6dB (right). The lines are the results of a linear fit of the data: in the noiseless case the slope *m* and the offset *q* were 1.39 and *−*1.62 *·* 10^*−*2^, respectively; in the noisy case, *m* = 1.62 and *q* = 8 *·* 10^*−*3^.

**Fig. 4:**
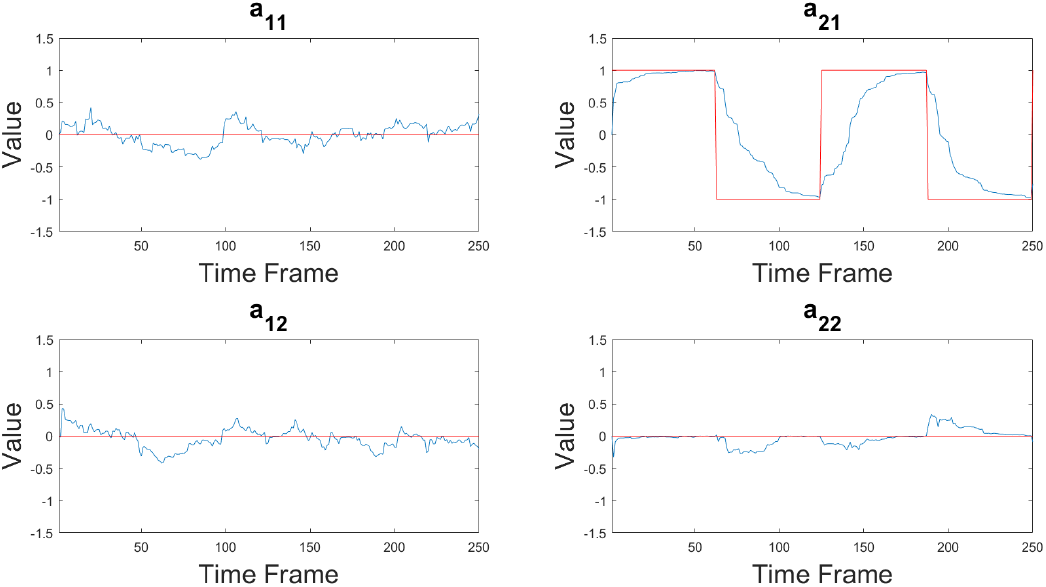
Time courses of the hidden parameters *a_ij_* in the case of a 2-node network with non-stationary coefficient *a*_21_ alternating between 1 and 1. Red lines represent the true values, while blue lines represent the estimates obtained by PF.

### B. Real fMRI Data

#### 1) Motor Network

Red lines in Fig. 5 represent average values of the *a_ij_* coefficients obtained on fMRI time series and the blue histogram represents the corresponding distribution of mean values of causal interactions for the permuted time series. Since temporal permutation suppresses the causal dependency between subsequent values, it was expected to observe zero-mean gaussianly distributed results, that is, no causal dependency. On the other hand, coefficients representing the causal relationship between two interacting brain areas should have values lying far away from the null distribution. This is what Fig. 5 shows, demonstrating that the particle filter effectively discriminates between unrelated and causally related time series.

**Fig. 5:**
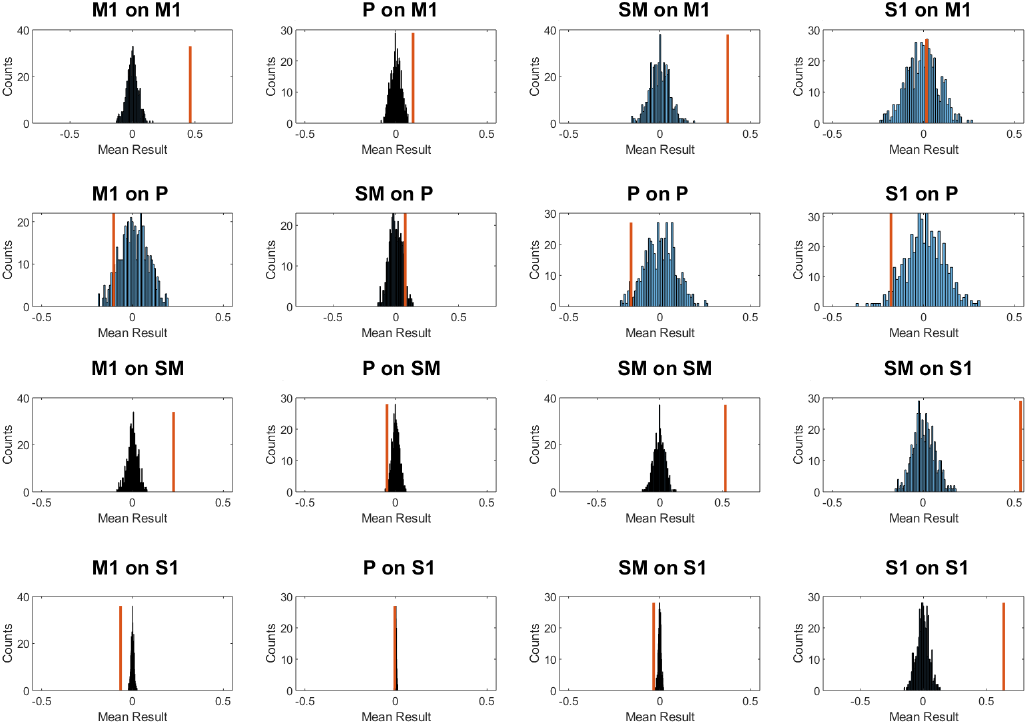
Mean results of causal interactions between real fMRI data computed through the particle filter: in blue, the histogram of mean values obtained on time-series randomly permuted in time, while the red line shows the corresponding mean value obtained from non-permuted time-series. Many of these values lie well outside of the null distribution, therefore their value reflects the effective mean causal interaction and it is not produced by chance.

The PF captured causal interactions between brain areas, which significantly correlated with a proxy measure of effective connectivity, that is, delayed correlation (DC) (p *<* 0.001, Pearson’s correlation coefficient *ρ* = 0.74, Fig. 6). In particular, in both subjects, the highest *a_ij_* coefficients in both PF and delayed correlation were those which represent the causal influence exerted by areas M1 and S1, in agreement with current knowledge of brain functioning during a sensory-motor task.

**Fig. 6:**
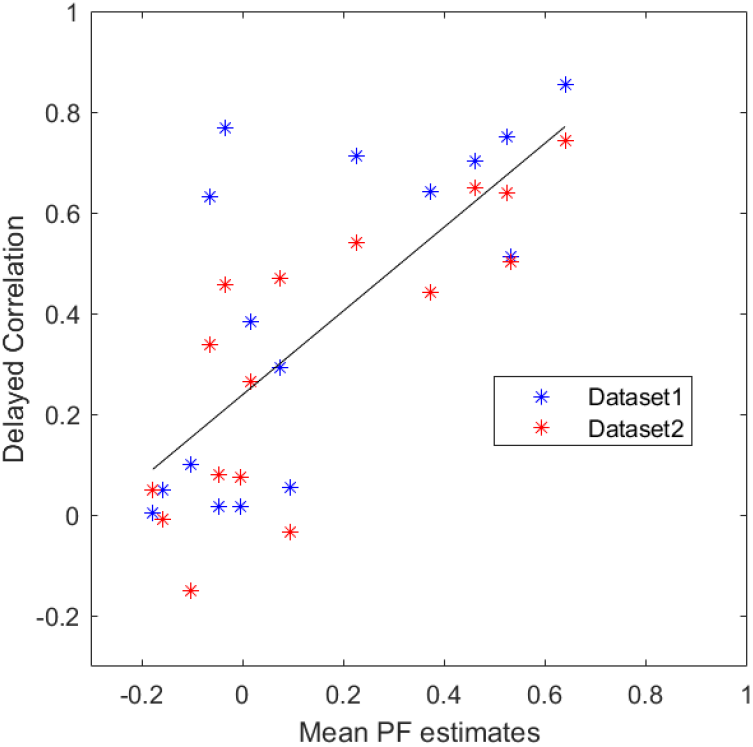
Scatter plot showing the relationship between mean PF estimates (horizontal axis) and delayed correlation (vertical axis) on the two sensory-motor experiments. On both results taken altogether the Pearson’s correlation coefficient *ρ* is 0.74, which corresponds to a statistically significant correlation with *p <* 0.001. Slope and offset of the linear fit were 0.83 and 0.24 respectively.

Fig. 7 exemplifies the temporal evolution of brain connectivity through three representative *a_ij_* coefficients in Subject 2. The top panel displays one representative coefficient involving the control ROI P, which is approximately 0. The two bottom panels demonstrate the expected reciprocal influence between M1 and SM areas. It is worth noting that the influences in the two directions, i.e. M1 over SM and vice versa, have different values, as a consequence of the adopted model that allows non-symmetric matrix of coefficients.

**Fig. 7:**
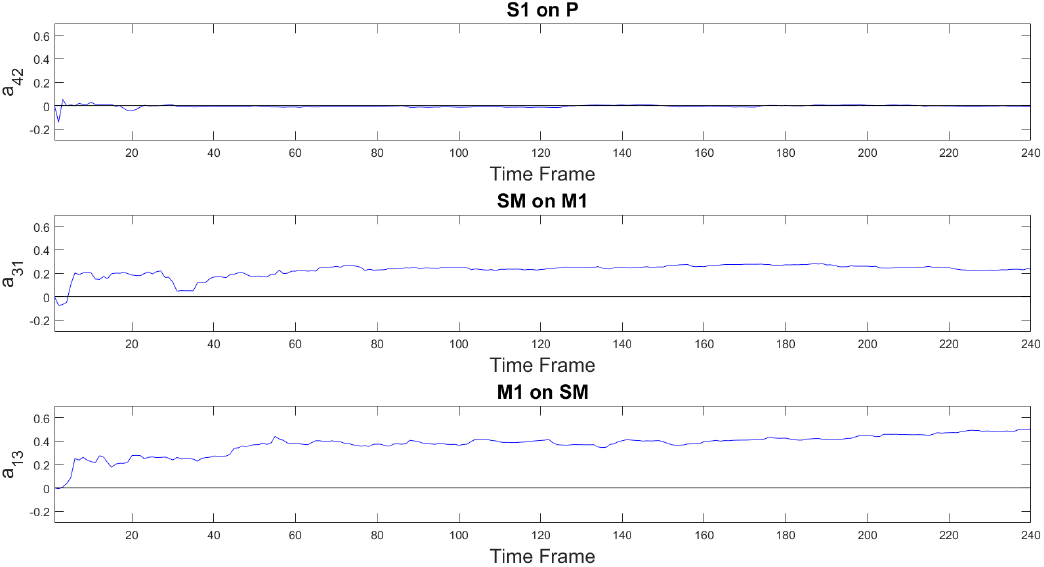
Plots in blue color show the PF-estimated time courses of three representative hidden parameters *a_ij_* in the case of a 4-node motor network, estimated in real fMRI data in one subject. Top panel depicts the coefficient describing the negligible causal effect exerted by area S1 over P. Central and bottom panels represent the causal effect exerted by the SM area over M1 and viceversa respectively.

#### 2) Visual Network

The PF detected causal dependencies between brain areas of the visual network which also correlated with the delayed correlation (Pearson’s *ρ* = 0.70 and *p <* 0.001, as shown in Fig. 8a). Interestingly, all four fMRI datasets showed statistically significant dynamic changes (*p <* 0.05) with a pattern following the underlying stimulation in the effective connectivity coefficient regarding the influence of MT on V1, which are known to take part in the processing of visual stimuli. As an example, Fig. 9 represents the statistically significant differences in the causal influence of MT on V1 in the presence or absence of the visual stimulus, observed consistently in all four fMRI datasets. Blue crosses show the quality parameter *Q* = *T/σ*, where *T* is the temporal length of the time series and *σ* is the standard deviation of the data, taken as a measure of noise. *Q* values indicate a possible explanation for inter-dataset variability: the particle filter performance are enhanced by the timeseries length (T) and reduced by noise (estimated as 1*/σ*). Not all connectivity coefficients share this same behavior, therefore other factors might influence the results. Fig. 10 represents one example of absence of detected causal relationship between area MT and the control region CTRL, and one example of non-symmetrical causality exerted by area V1 over MT and viceversa.

**Fig. 8:**
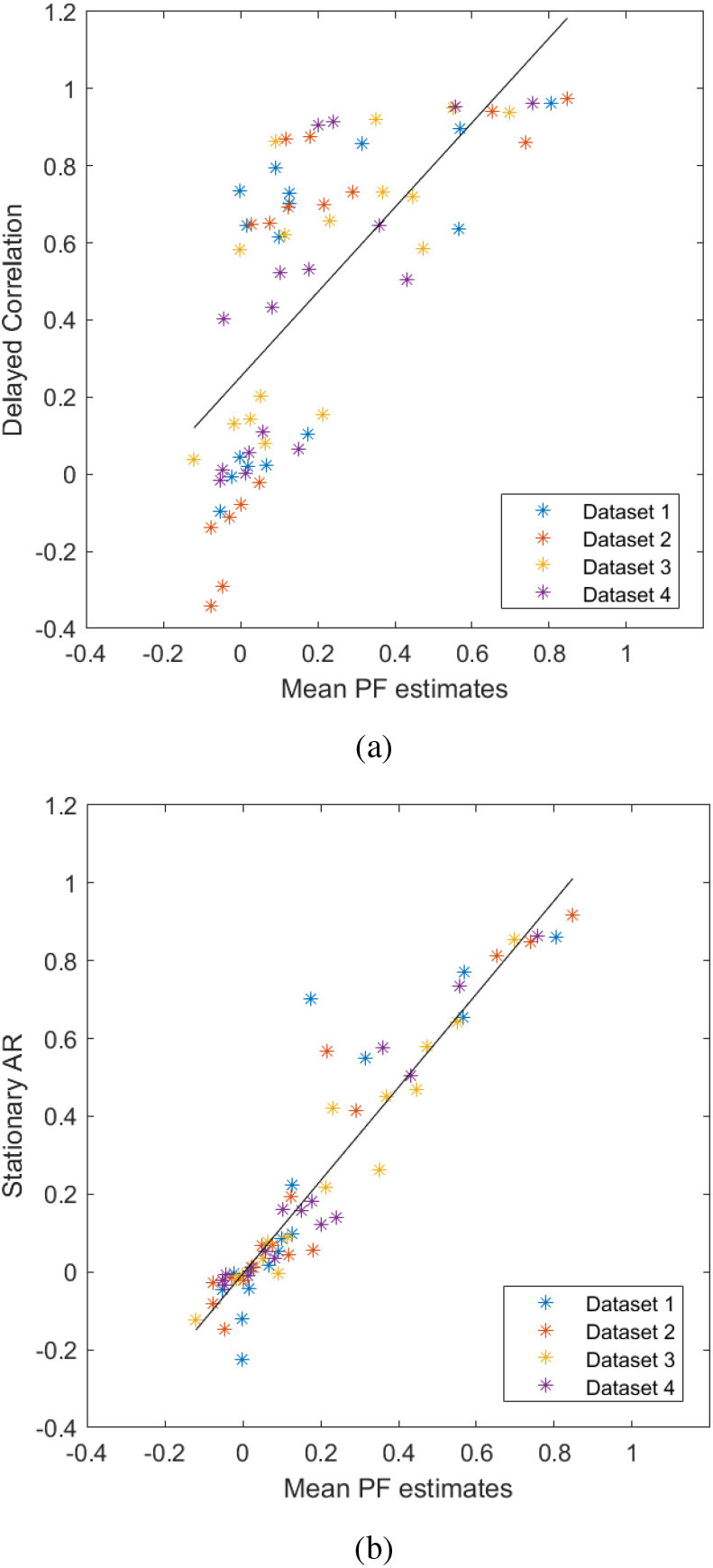
(a) Scatter plot between mean PF estimates (horizontal axis) and delayed correlation (vertical axis) on the four visual experiments fMRI data. A *p <* 0.001 was obtained with a Pearson’s correlation coefficient *ρ* = 0.70. The black line is the result of a linear fit. Slope and offset of the linear fit were 1.10 and 0.25 respectively. (b) Scatter plot between average PF estimates (horizontal axis) and stationary AR coefficients (vertical axis). The resulting Pearson’s correlation coefficient *ρ* was 0.94, with a *p <* 0.001. The black line is the result of a linear fit, with slope and offset 1.2 and −0.004 respectively.

**Fig. 9:**
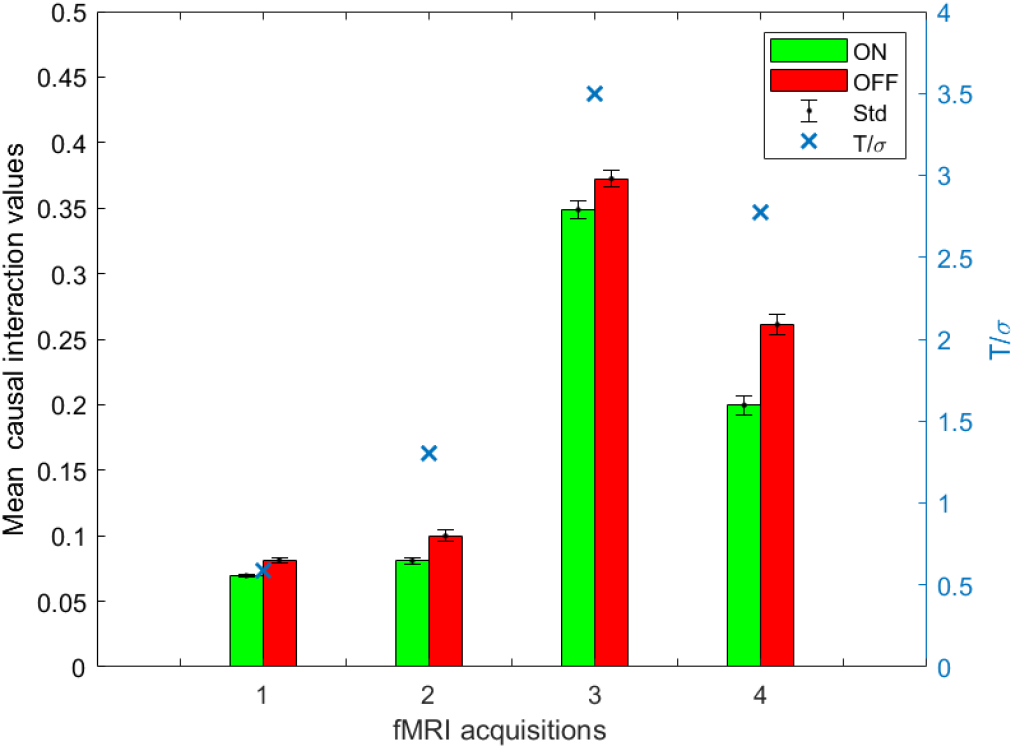
Comparison between presence (green bar) and absence (red bar) of visual stimulation for the mean *a_ij_* coefficient representing the causal influence of MT on V1 in the four different datasets. Bars indicate the mean values and the standard deviation of the mean. Higher values of the coefficients are obtained in data sets with better quality parameter *Q*, as shown by blue crosses.

**Fig. 10:**
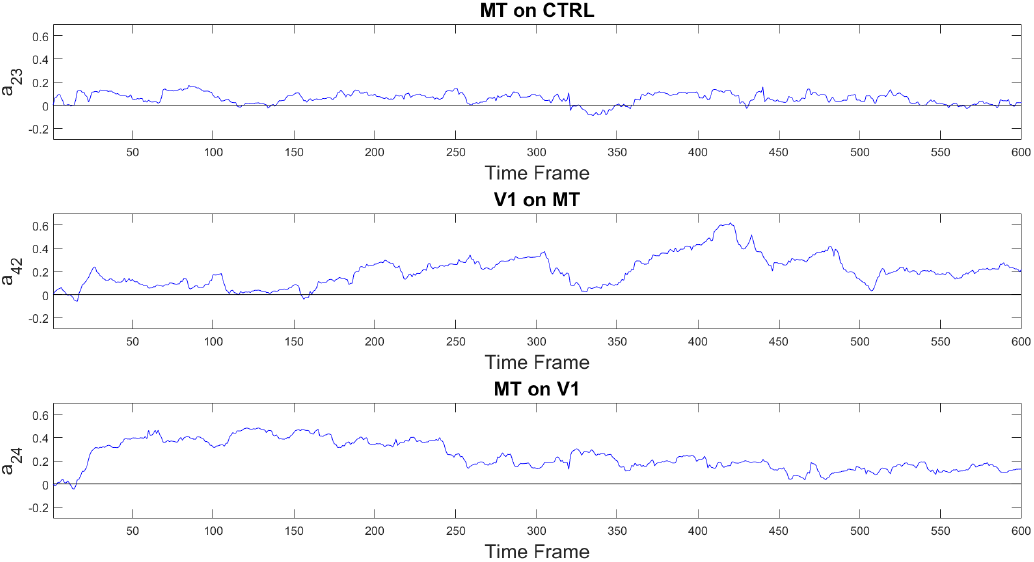
Plots in blue color show the PF-estimated time courses of three representative hidden parameters *a_ij_* in the case of the 4-node visual network, estimated in real fMRI data in one subject. Top panel depicts the coefficient describing the negligible causal effect exerted by area MT over the control area. Central and bottom panels represent the non-symmetrical causal effect exerted by the V1 area over MT and viceversa respectively.

Fig. 8b shows the comparison between results of the PF and stationary AR coefficients (Pearson’s *ρ* = 0.94 and *p <* 0.001). An excellent agreement was found between the two methods. In addition, changes through time of the AR coefficient were searched through a sliding-window approach. While in this way some coefficients were found to vary, the results were not consistent between the four datasets.

Fig. 11 shows the relationship between coefficients obtained by PF and DC with explicit reference to the causal influence they are referred to. We searched for the appropriate clustering of the data in Fig. 11. The number of clusters was chosen as the knee of the number-of-clusters vs distance-to-every-centroid curve. The optimal number was 3 and each cluster’s centroid was plotted as a black cross in Fig. 11. All coefficients representing the causal interactions involving the CTRL region (black symbols) belong to cluster A, that is, small values of DC and mean PF estimates, except for the self-causality terms, which all lay in the vicinity of the C centroid. Every nondiagonal coefficient involving ROIs LGN, MT and V1 belong to the cluster B. In particular, in Fig. 11, the horizontal red line and vertical green line intersect in the point equally distant to the centroids A and B. All values representing the causal influence of MT on V1 (which vary following the stimulation pattern, represented by highlighted red diamonds in the figure) and that of V1 on MT (green stars) lay right to the green line and above the red line.

**Fig. 11:**
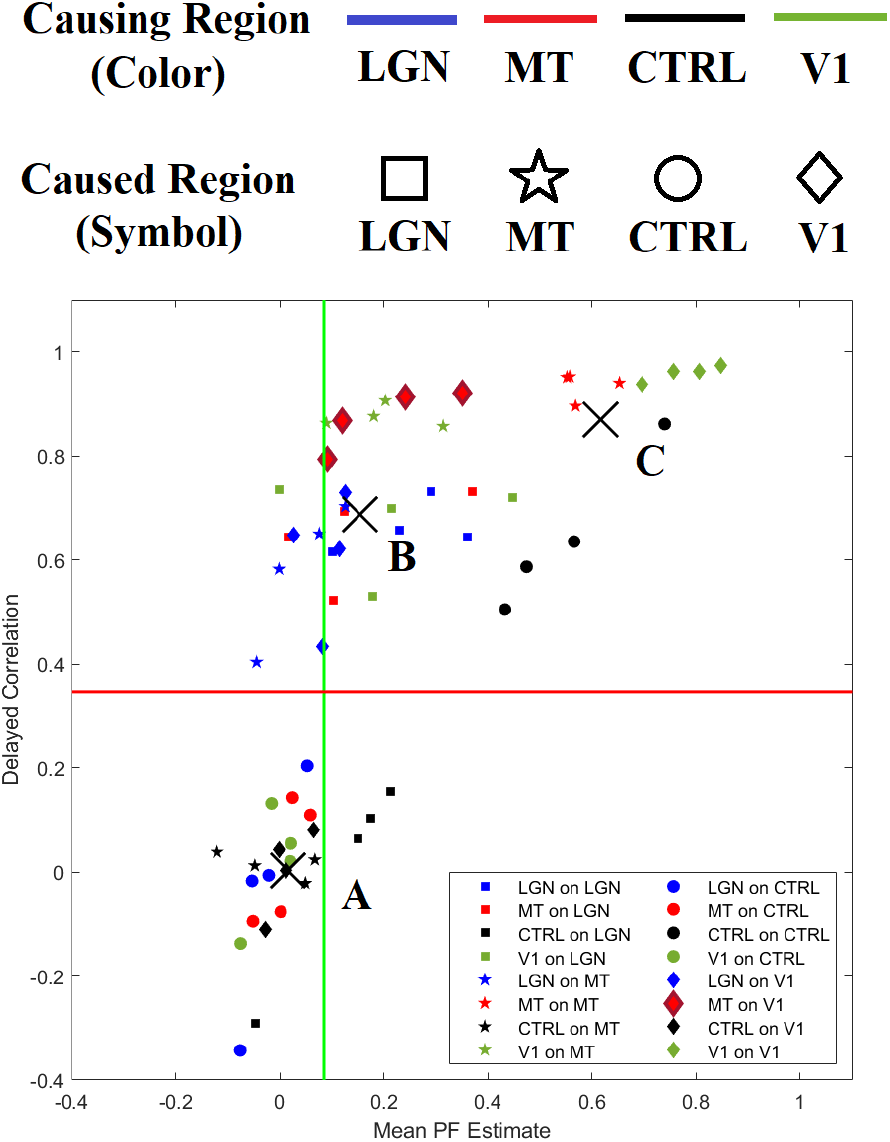
Fig. 8a with ROI-based encoding, explained in the legend above the figure: same color refer to the same *causing* ROI and same symbols to same *caused* ROI. Black crosses are clusters centroids found through the Matlab function *kmeans()*, called A, B and C as in figure. The horizontal red line and vertical green line intersect in the point equally distant from centroids A and B.

## IV. DISCUSSION

Other implementations of SMC algorithms were previously proposed to investigate brain connectivity in fMRI data. Murray and Storkey [28] proposed a forward-backward Particle Filter using, as observation equation, a stochastic extension of the balloon model, which was proposed to describe the haemodynamics that follow brain activity [29]. In their study, the hidden parameters of the model resulted approximately constant, probably as a consequence to the complexity of the model itself [30], [31].

In a different study, Ahmad et al. [32] adopted a symmetric, linear, first-order, time-varying Autoregressive (TVAR) model and used a Rao-Blackwellized PF to estimate the temporal relationships among fMRI time-series representing four brain regions during resting state. The assumption of symmetry in coupling coefficients, that is *a_ij_* = *a_ji_*, reduced model complexity, but did not permit to infer neither the directionality of the network nor any possibly asymmetric cause/effect interaction between brain areas. Therefore their approach cannot be used to investigate effective connectivity. Also, the results were not benchmarked with the outcome results from different analyses and the resting-state paradigm did not allow any analysis on the temporal evolution of the results.

In our implementation on fMRI data, the time-averaged PF estimates were in agreement with a proxy measure of causality, that is, delayed correlation. Part of the mismatch between the proposed method and delayed correlation could be explained by the fact that the PF algorithm studies the network as a whole and produces estimates of *a_ij_* coefficients that update at every time instant, while delayed correlation is a measure of pairwise causality that does not take into account possible nonstationarities and spurious cause-effect relationships mediated by other nodes of the network.

In the first experimental setup involving the motor network, statistically significant changes in connectivity were not found, but the poor temporal resolution of the data (2*s*) may have prevented the detection of these changes.

On the contrary, in our study of the visual network statistically significant changes in connectivity were found in four different datasets with a pattern following the underlying stimulation, without requiring any previous knowledge on the actual stimulation paradigm during the estimation process. These variations interested the influence of MT on V1 which is consistent with our understanding of cortical processing in the early visual cortex [24]. Good agreement was found between PF results and an AR coefficient evaluation method, even if the latter failed in detecting variations through time in a slidingwindow fashion. This result suggests that this second method might require more data, i.e. longer exams, to achieve the sensitivity obtained with the PF. Furthermore, while the PF is “blind,” sliding window analysis require the inclusion of information regarding the stimulation timing, which in some cases may not be available.

Mean causal interaction values were found to vary between different datasets. As Fig. 9 shows, in some cases these differences can be explained through the simultaneous contribution of noise and temporal length of the data, i.e. the quality parameter *Q* = *T/σ*. The brain haemodynamic responses may also be involved, which vary not only among subjects but also between different areas in the same subject [33], which was not taken into account in this study.

## V. CONCLUSIONS

We used Particle Filter to test and identify the timevarying brain connectivity as evidenced in fMRI images. Our experiments confirmed the hypothesis of time-varying brain connectivity pattern and gave evidence for non-symmetric connectivity. It was possible to detect statistically significant changes in cortical cause-effect relationships correlated with the underlying task-rest pattern during the fMRI acquisition.

Future studies should test the performance of the proposed algorithm in fMRI experiments with higher time resolution, namely *<* 0.8*s*, and they should aim to unveil possibly asymmetric changes in effective connectivity among brain regions. Also, to minimize the impact of vascular dynamics and highlight neural ones, future studies should use more sophisticated experimental designs that enable a better control over the non-uniformity of brain haemodynamics across different areas [33]–[35].

As suggested by Bugallo and Djuric [36], the PF can be improved by a parallel implementation when dealing with complex system, such as the brain. Ordinary brain connectivity analysis represents ROIs as time series obtained by averaging signals originating from more voxels of that region. This helps improving the Signal-to-Noise Ratio (SNR) but assumes some wide-scale connectivity features. Because of this wide-scale connectivity assumption, more reliable results may be achieved with a parallel combination of Particle Filters carried over single-voxels time series.

Our results, together with the possibility to refine the methodology, suggest that the proposed computational method, the Particle Filter, can be capable to infer timevarying effective connectivity on acquisitions with acceptable scan duration, without the need of any constraint or previous knowledge about the examined network or timing of the underlying brain processes.

## VI. ACKNOWLEDGMENTS

The authors thank Dr Sergiy Ancherbak for having shared with them his particle filtering code for time-dependent gene network modeling. This work has been partially supported by grants “RC 2018-2020” and “5 per mille” to IRCCS Fondazione Stella Maris, funded by the Italian Ministry of Health.

